# FastConformation: A Standalone ML-Based Toolkit for Modeling and Analyzing Protein Conformational Ensembles at Scale

**DOI:** 10.1101/2025.05.09.653048

**Authors:** Flavia Maria Galeazzi, Gabriel Monteiro da Silva, Pablo Arantes, Iz Varghese, Ananya Shukla, Brenda M. Rubenstein

## Abstract

Deep learning approaches like AlphaFold 2 (AF2) have revolutionized structural biology by accurately predicting the ground state structures of proteins. Recently, clustering and subsampling techniques that manipulate multiple sequence alignment (MSA) inputs into AlphaFold to generate conformational ensembles of proteins have also been proposed. Although many of these techniques have been made open source, they often require integrating multiple packages and can be challenging for researchers who have a limited programming background to employ. This is especially true when researchers are interested in subsampling to produce predictions of protein conformational ensembles, which require multiple computational steps. This manuscript introduces FastConformation, a Python-based application that integrates MSA generation, structure prediction via AF2, and interactive analysis of protein conformations and their distributions, all in one place. FastConformation is accessible through a user-friendly GUI suitable for non-programmers, allowing users to iteratively refine subsampling parameters based on their analyses to achieve diverse conformational ensembles. Starting from an amino acid sequence, users can make protein conformation predictions and analyze results in just a few hours on their local machines, which is significantly faster than traditional molecular dynamics (MD) simulations. Uniquely, by leveraging the subsampling of MSAs, our tool enables the generation of alternative protein conformations. We demonstrate the utility of FastConformation on proteins including the Abl1 kinase, LAT1 transporter, and CCR5 receptor, showcasing its ability to predict and analyze the protein conformational ensembles and effects of mutations on a variety of proteins. This tool enables a wide range of high-throughput applications in protein biochemistry, drug discovery, and protein engineering.

## 1 Introduction

AlphaFold 2 (AF2), a deep learning-based approach for predicting the 3D structures of proteins, has revolutionized structural biology due to its ability to rapidly predict the ground state structures of proteins with high accuracy. ^1^ AF2 employs a deep neural network architecture to generate its predictions of three-dimensional protein structures. AF2 was trained on Uniprot, which contains co-evolutionary data in the form of multiple sequence alignments (MSAs), as well as the Protein Data Bank (PDB), which contains solved protein structures. When an amino acid sequence is input, AlphaFold 2 (AF2) generates multiple sequence alignments (MSAs) to compare the target sequence to a vast database of known sequences. The neural network then processes these MSAs, extracting co-evolutionary features at each layer. By analyzing these features, AF2 predicts the distances and orientations between amino acids, ultimately constructing a three-dimensional model of the protein. The initial model is then refined to adhere to physical and chemical constraints arising from AF2’s training on the PDB. AF2 differs from prior attempts at protein-structure prediction because it infers patterns of interactions among amino acid sequences in the MSAs rather than predicting and/or sampling an energy function. This type of evolutionary coupling works because amino acids co-evolve together to perform different functions; therefore, it follows that their three-dimensional arrangement must be encoded in their sequence.^2^ AlphaFold’s ability to rapidly predict protein structures has had significant implications for understanding protein function,^3^ analyzing protein interactions,^4^ and furthering the development of new therapeutics.^5,6^

Despite these clear impacts, AlphaFold2 is still limited in its capacity for predicting the alternative conformations (or states) of proteins, which is an important consideration for structural biology and drug discovery. When used to predict the structure of a sequence with standard parameters, AlphaFold2 will mostly predict very similar structures, no matter how many times it is run, even for proteins that are known to fold-switch and occupy different conformations at room temperature.^7,8^ This overwhelming preference for low-energy states is neither accidental nor a detriment to AlphaFold’s accuracy, as its inference model was designed using low-energy structures deposited in the Protein Data Bank (PDB), generated from X-ray crystallography or cryogenic electronic microscopy experiments. Since AlphaFold’s training data is overwhelmingly composed of low-energy states, it follows that it excels at predicting the same. However, understanding proteins’ conformational ensembles has significant impacts in structural biology, health, and drug discovery. Determining the conformational ensemble of proteins and how mutations affect these conformations is important for understanding disease mechanisms and the consequent development of therapeutics.^9^ For example, in cancer, mutations in the Ras protein can shift its conformational equilibrium, leading to persistent activation of signaling pathways that drive uncontrolled cell growth.^10^ Understanding these conformational changes provides insight into how to design targeted inhibitors that can lock the protein in an inactive state, slowing disease progression. Exploring the conformational landscapes of proteins is currently challenging, and not difficult to scale, relying on complex experiments such as nuclear magnetic resonance or small-angle X-ray scattering, or computationally-expensive, physics-based molecular dynamics simulations (MD).

Ref. 11 demonstrated that restricting the depth of the MSA AF2 uses as an input biases AlphaFold towards predicting alternative protein conformations by turning the attention of the network to different parts of the MSA, which allows it to find alternative conformations based on other co-evolved residues. Different methods for generating conformational ensembles have also been explored, including masking,^11^ clustering,^12^ or randomly subsampling the input multiple sequence alignment (MSA). ^13^ All of these methods have been shown to increase the probability of predicting physiologically-relevant alternative conformations that are not yet deposited in the PDB, which can then be used as seeds for MD simulations.^14^ Having access to ensemble predictions at scale would thus be as revolutionary for these purposes as AlphaFold’s ground state prediction accuracy was.

Using subsampling, our group has shown that AF2 can be used to predict the effects of single and double point mutations on the conformational distributions of the Abl1 tyrosine kinase core and a wide range of other proteins. ^14^ We have also shown that subsampling can be used to predict changes in the conformational ensemble of granulocyte-macrophage colony-stimulating factor (GMCSF), a protein with minimal known homology, in response to point mutations. Subsampled GMCSF predictions strongly correlated with experimentally-determined NMR results, further demonstrating subsampled AF2’s remarkable capacity to decode signals pertaining to conformational changes even when sequence data is scarce.^14^

These distributions are crucial for predicting protein function and understanding biological pathways. By comparing AF2 predictions with NMR results, we can validate and refine NMR models, enhancing their accuracy.^14^ Furthermore, the ability to predict conformational changes in mutants enables clinical comparisons, which can be instrumental in drug discovery and personalized medicine. Additionally, these conformational distributions can serve as initial structures for molecular dynamics (MD) simulations. ^15^ Altogether, these results highlight the strong, yet untapped potential of AF2 for predicting the conformational ensembles of proteins, which will have substantial impacts in the fields of biophysics and drug discovery.

Several efforts, including ColabFold^16^ and OpenFold,^17^ have made AF2 more accessible, allowing users to predict structures using a modified AF2 protocol run on an open-access server. For instance, ColabFold uses AlphaFold2 internally and can predict protein structures faster by employing MMseqs2 for MSA generation. ColabFold can be run on Google Colaboratory (Colab) or a local machine with fewer computational requirements. It is also available via a command-line interface or a Python package hosted on PyPI (Python Package Index). However, there are several steps need to adapt the pipeline for conformational ensemble prediction. The user must optimize the MSA generation method and prediction parameters, extract relevant information from the predictions, and analyze the results. This opens up the possibility of combining these steps into one package that allows users to generate the input MSA, run the predictions, and visualize the results in a way that is fast, interactive, and accessible to the scientific end-user.

FastConformation is a Python-based application that offers these functionalities all in one place. Starting from an amino acid sequence, the user can predict different protein conformations and analyze the results in only a few hours on their local machine. FastConformation can be executed without any programming knowledge via a Graphical User Interface (GUI). The GUI combines different MSA generation methods (MMSeqS2^18^ and JackHMMeR^19^), fast protein structure prediction internally run by AlphaFold2 through ColabFold, and interactive analysis of the AF2 output. Our interactive analysis features allow the user to identify the optimal subsampling parameters to predict multiple states of their protein in a quick, interactive, and accessible way without any specialized computer knowledge. We demonstrate the facility and applicability of our tool by testing it on three different example systems: Abl1 kinase,^20^ LAT1,^21^ and CCR5.^22^ FastConformation makes protein structure prediction methods more user-friendly and may be adapted for a variety of important applications in protein biochemistry, drug discovery, and protein engineering.

## 2 Methods

### 2.1 Overview of AlphaFold 2

AlphaFold 2 (AF2)^1^ represents a groundbreaking advance in protein structure prediction, leveraging deep learning to predict the ground state structures of proteins with near-experimental accuracy.^23^ To do so, the algorithm integrates multiple sequence alignments (MSAs) and structural templates to predict 3D protein structures from amino acid sequences. AF2’s architecture comprises two key components: the Evoformer module and the Structure module. The Evoformer processes MSAs and structural templates, capturing long-range interactions and co-evolutionary patterns using sophisticated attention mechanisms such as axial attention.^1^ A pair representation matrix further encodes inter-residue relationships. The Structure module translates these refined features into 3D atomic coordinates. AF2 employs a custom loss function to ensure its predictions conform to physical and chemical constraints. During inference, MSAs are generated from the target sequence and then processed through the Evoformer and Structure modules. The output is refined through iterative cycles of recycling and optimization, significantly enhancing the algorithm’s accuracy. Trained on diverse datasets, including the Protein Data Bank^24^ and metagenomic sequences, AF2 demonstrates predictive accuracy on par with experimental methods for many ground-state protein structures.^25^

### 2.2 AlphaFold 2 Subsampling for Conformational Ensembles

Although AF2 excels at predicting ground state structures, its architecture is less suited for capturing alternative conformations, which are critical for understanding protein function and dynamics. Alternative conformations play essential roles in biological processes, such as the activation and inactivation of proteins, and influence protein interactions, including drug binding.^26^ Thus, predicting conformational ensembles can vastly improve drug discovery by providing insights into different structural states.^27^

AF2’s ability to interpret sequence data can be extended to predict conformational variability. The underlying hypothesis is that distinct conformational states are encoded within protein sequences. By subsampling the input MSA, it is possible to modulate co-evolutionary signals across structural domains, promoting the prediction of alternative conformations.^14^

In a previous study,^14^ we introduced a subsampling strategy to overcome AF2’s limitations in predicting conformational diversity. This method involves modulating co-evolutionary signals within MSAs by clustering sequences using a Hamming distance. By selecting different subsets of sequences, modulated by two key parameters, ‘*max_seq*’ (number of cluster centers) and ‘*extra_seq*’ (sequence sample size per cluster), our ‘subsampled’ AF2 generates a range of conformational states.

In this work, we focus on FastConformation, a user-friendly tool that automates this subsampling approach, enabling researchers to easily apply it to their systems of interest. FastConformation provides a flexible interface for configuring parameters and running ensemble predictions in a high-throughput manner. Users can explore the structural diversity of their proteins without needing to delve into the technical complexities of MSA manipulation, making this powerful method accessible to a broader audience.

The variance of the ensembles depends on the chosen parameters: lower values of ‘*max_seq*’ and ‘*extra_seq*’ yield greater structural diversity, while higher values result in more conservative predictions. To determine the optimal parameters, users must evaluate the balance between ensemble diversity and prediction confidence. This involves examining root mean square fluctuation (RMSF) plots to identify highly flexible regions, comparing pLDDT scores to ensure high-confidence predictions, and assessing root mean square displacement (RMSD) distributions to verify that alternative conformations are physiologically relevant. By iteratively adjusting ‘*max_seq*’ and ‘*extra_seq*,’ users can fine-tune the subsampling to obtain the most informative structural ensemble for their protein system. FastConformation further enhances statistical power by enabling multiple rounds of prediction with different random seeds. This iterative approach has successfully predicted diverse conformational states across several systems,^14^ making FastConformation a powerful tool for high-throughput ensemble prediction.

### 2.3 FastConformation GUI

The FastConformation graphical user interface (GUI) provides an intuitive workspace for managing three core workflows: MSA generation (Build MSA), ensemble prediction (Make Predictions), and analysis (Analysis). The menu bar offers options to display logs and monitor task progress. While MSA generation and ensemble prediction operate in the background, the analysis mode provides real-time, interactive visualizations. The GUI can be installed by running the install.sh script found on the GitHub page.

Upon launch, the home window offers users the option to initiate new tasks (Figure 1). Selecting “Submit New Job” opens a configuration window, where users choose between MSA generation, structure prediction, or analysis. Additional settings allow customization of parameters and computational engines. Tasks are executed locally and results are saved to a user-defined output directory (Figure 2).

**Figure 1:**
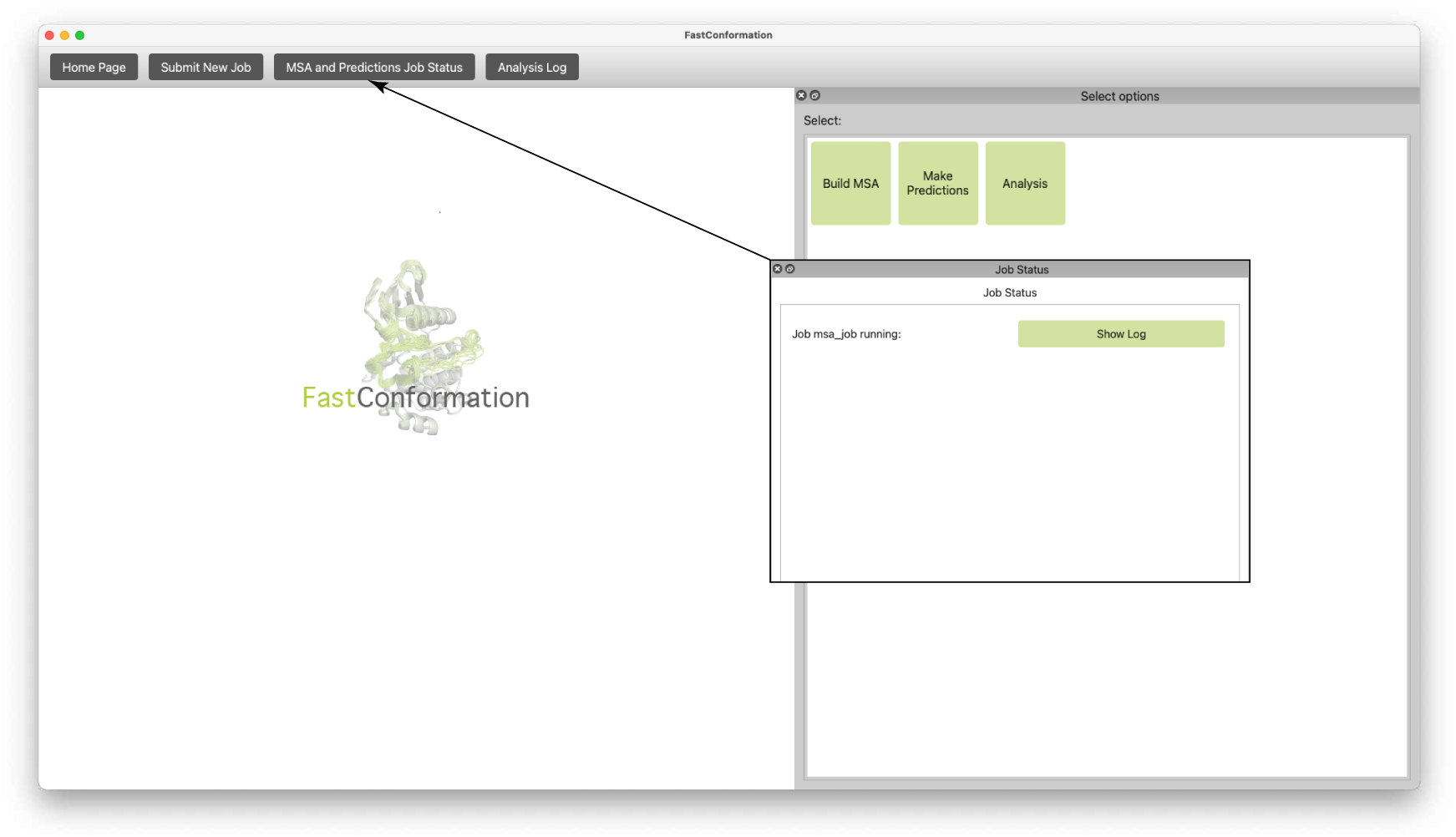
Home page of the GUI. The user is able to navigate to the three main menus of the GUI: MSA Generation (Build MSA), ColabFold Structural Ensemble Prediction using the generated MSA (Make Predictions), and output analysis (Analysis). The user can monitor the status of the MSA generation and Structure Prediction jobs and the progress of the Analysis by selecting the appropriate options on the top bar.

**Figure 2:**
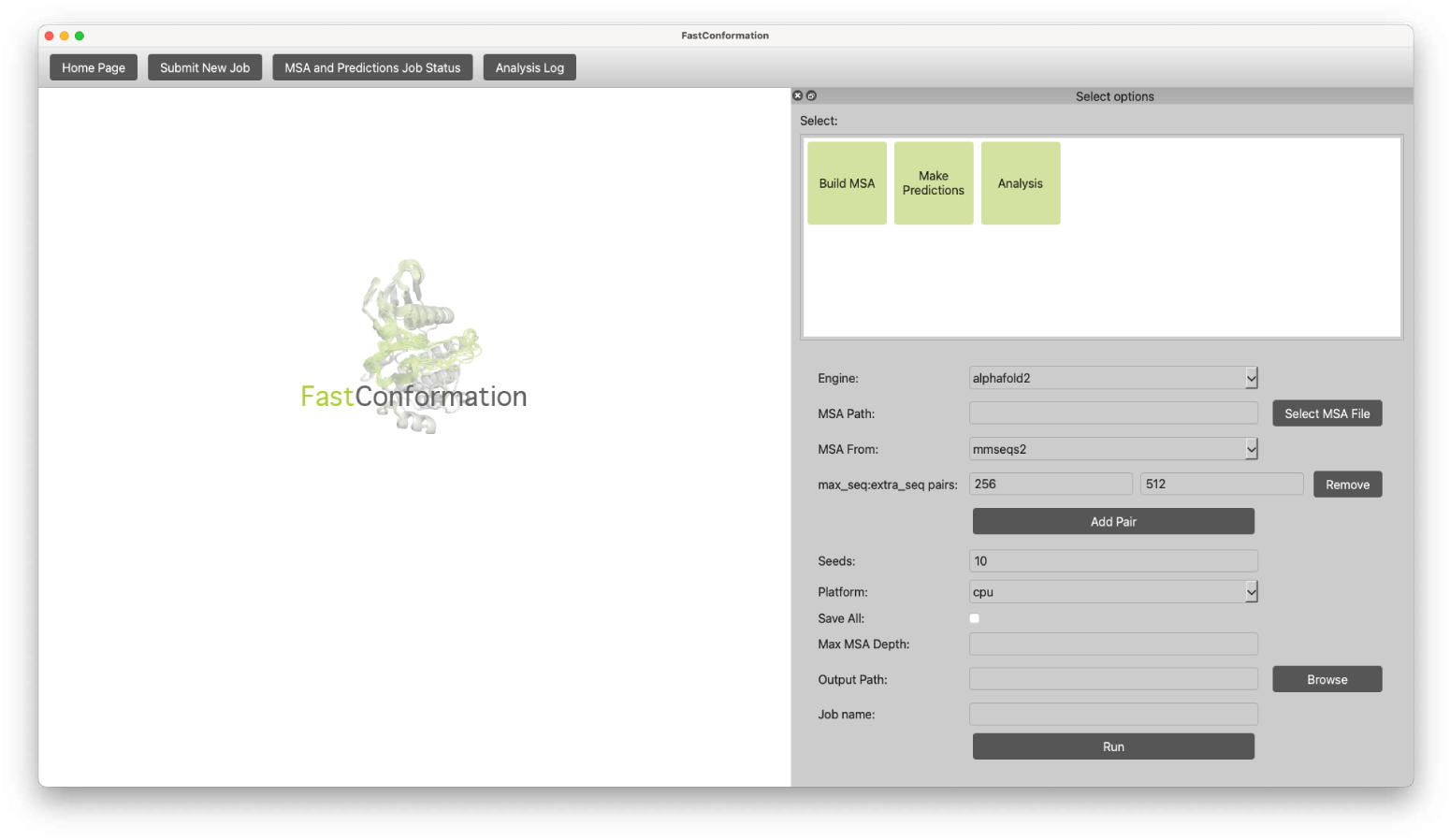
To submit an Ensemble Prediction job, the user navigates to the Make Predictions menu. Here, users can input the path to the previously-generated MSA, the *max_seq*:*extra_seq* parameters for subsampling, the number of seeds to repeat the predictions (Seeds), whether to run the code on a CPU or GPU (Platform), the option to save the predictions to a pickle file (Save All), the Max MSA Depth, and the output path for saving predictions. The user can then monitor the status of the job by selecting the “MSA and Predictions Job Status” option on the top bar.

In analysis mode, users load prediction results and configure visualization settings. The GUI displays plots interactively in the left panel, while task progress is shown in the log panel at the top (Figure 1). This centralized, user-friendly interface streamlines complex workflows, enabling efficient exploration of protein conformational ensembles.

### 2.4 FastConformation Implementation

The architecture of FastConformation consists of a Python-based core that acts as a graphical front-end for user interaction and communicates with ColabFold to predict protein structures from amino acid sequences via the Python subprocess module. FastConformation is available as a PyPI package for the Linux and Mac Operating Systems. For Windows 10 OS, FastConformation must be run on a WSL instance. FastConformation is written in Python 3.12 using the PyQt5 GUI framework. Third-party libraries may be obtained through PyPI or using a package manager and are included in a requirements file. Our application runs on the Central Processing Unit (CPU). For the analysis module, Graphics Processing Unit (GPU) acceleration is supported through the MDAnalysis and OpenMM packages. We calculate the TM-Score to assess protein structure similarity by running the original program through the Python subprocess module.

FastConformation can be installed by running a single script, which sets up the conda environment, installs MicroMamba if it is not already installed, and installs all dependencies. At the end of the installation, we also run a script to test the installation and configuration of the environment.

Installation instructions and extensive documentation are available on the GitHub page and ReadTheDocs.^28^ We also include tutorial videos which aim to guide the user through navigating the user interface, available on our GitHub page https://github.com/GMdSilva/FastConformation. To get started, we recommend running the sample input sequence provided and replicating the sample outputs to verify the installation.

During each step of the MSA generation, structure prediction, and analysis, the files are saved to an output directory provided by the user and displayed in the FastConformation GUI. The user can specify different sets of subsampling parameters run as a single job on the same protein sequence. Similarly, the user can analyze and compare the results of different subsampling parameters by plotting them on the same graph or on different graphs in the same widget window (Figure 3). This allows users to quickly and interactively assess the best set of subsampling parameters for their system.

**Figure 3:**
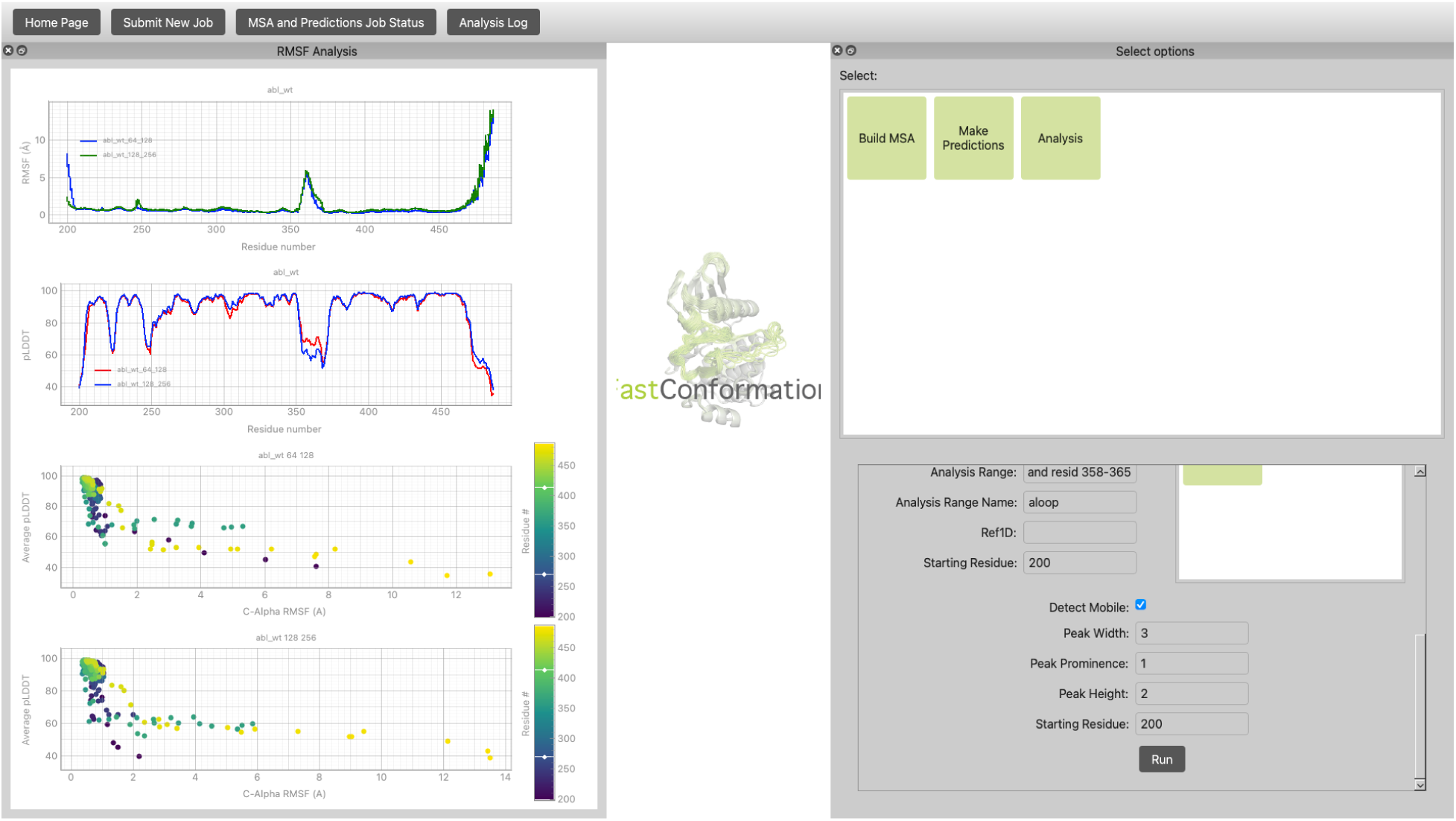
To analyze the generated structural ensembles predictions, the user navigates to the Analysis menu, selects the desired analysis method and the relevant parameters. After clicking “Run”, the plots will be shown on the left. Each plot is interactive and the user is also able to export them. Different *max_seq*:*extra_seq* pairs are plotted as different lines on the same plots, allowing users to iteratively refine their choices. Users should identify parameter sets that maximize structural diversity while maintaining realistic confidence scores (high pLDDT and reasonable RMSD values).

### 2.5 FastConformation MSA Generation and Subsampling

In MSA subsampling methods, an MSA used as input to AlphaFold2’s EvoFormer stack is stochastically subsampled by first selecting *N* sequences as cluster centers (*N* is defined by the *max_seq* parameter). Importantly, the target sequence is always selected as a cluster center. This new MSA is then piped to the EvoFormer’s row/column attention track, from which most of the coevolutional data is distilled.

The subsampled MSA is then constructed by adding *M* sequences, where *M* is defined by the *extra_seq* parameter, which were not previously selected as cluster centers around the *N* cluster centers. This second MSA also affects the EvoFormer stack without going through the row/column attention track.

The *max_seq* and *extra_seq* parameters employed have a strong influence on the diversity of the structures predicted by AF2. Users must experiment with different parameter combinations to tune the diversity of the predicted ensemble for their system. Typically, lower values of *max_seq* and *extra_seq* promote greater diversity, but too large of a reduction may lead to unphysical structures. By analyzing RMSF plots for flexibility patterns, RMSDs for cluster distributions, and pLDDT values for structure confidence, users can iteratively refine their choices to produce a well-balanced conformational ensemble. In what follows, we describe the analysis tools FastConformation provides to facilitate parameter choices.

### 2.6 FastConformation Analysis Tools

A common problem when analyzing protein conformational ensembles is identifying mobile structural elements (and measuring the amplitude of that mobility). Thus, we have developed the rmsf plddt module of FastConformation to streamline and facilitate the identification of mobile elements. In this module, high-mobility elements in a collection of .pdb files defining an ensemble are identified via the RMSF of *α*-carbon (CA) positions metric. This metric identifies regions of high mobility within the protein. Peaks in the RMSF plot highlight areas with significant structural flexibility. The RMSF is calculated after aligning each element in the ensemble to a reference formed by the average positions of each CA in the ensemble. The resulting RMSF values are distributed starting from the first CA position, and continuous peaks of user-specified width, height, and prominence are automatically detected, representing residue ranges of enhanced flexibility. The CA ranges detected through the above are then saved to a report file, along with accompanying plots illustrating the detection process.

Beyond RMSF values and peak-calling results, we also include line plots representing per-CA average pLDDT metrics for each ensemble, as well as 2D scatterplots representing the pLDDT/RMSF correlation for each CA position. This information is presented to further facilitate the selection of mobile segments, as highly-mobile CAs tend to have relatively large RMSFs and relatively small pLDDTs (that is, AlphaFold2 is less confident regarding their positions).

The RMSD function in the GUI calculates the deviation between the predicted structures and a reference structure. This analysis is particularly useful for evaluating the effects of mutations or sequence variations on protein dynamics. By comparing the RMSD values across different predictions, users can assess how specific changes in the sequence impact the overall protein conformation. This function is also employed to detect unphysical structures, indicated by large RMSD values in regions expected to be stable. We similarly use a peak detection algorithm in the sci-kit package to identify and highlight noteworthy peaks in the RMSD plots.

The template modeling score (TM-score) ^29^ function in the GUI is used to compare the predicted structures against reference structures to assess their structural similarity. This metric provides a quantitative measure of the accuracy of the predicted protein models. By calculating the TM-score for the ensemble of predictions, users can identify the most reliable models and understand the range of structural variations captured by the subsampling approach.

The two-dimensional RMSD and TM-score functions in the GUI extend the analysis to pairwise comparisons across all predicted structures. These plots provide a comprehensive view of the structural relationships within the ensemble, facilitating the identification of clusters of similar conformations. Such an analysis helps distinguish among different conformations and facilitates understanding the overall structural diversity within the ensemble.

A similar method can also be applied to determine the effects of changes in the sequence on protein dynamics. By calculating the RMSD between a mutated and reference structure, it is possible to gain insights into how that change in the sequence affects protein dynamics in target regions of the protein. For example, if a specific combination of subsampling parameters results in high variation in predicted distance probabilities in regions of the protein that are predicted to be static by other parameter sets or where dynamics are not expected based on prior knowledge, this may indicate the presence of unfolded or unphysical structures.

In the following, we illustrate the functionality of FastConformation on three example proteins of clinical significance.

## 3 Illustrative Examples of FastConformation Applications

### 3.1 Conformations of the Abl1 Wild Type Protein and Its Mutants

Abl1 is a non-receptor tyrosine kinase involved in a variety of cellular processes, including cell differentiation, division, and response to stress. One of its key functions is its ability to phosphorylate substrates, a process that can be tightly regulated by its own phosphorylation status, which can either activate or inactivate its kinase activity.^30,31^ The structure of Abl1 consists of conserved N- and C-lobes, with key structural elements involved in its dynamics, including the activation loop (A-loop), the Asp-Phe-Gly (DFG) motif, the glycine-rich loop (G-loop), and the *α*C-helix (Figure 4).^32^ One of the primary mechanisms for deactivating Abl1 is the DFG flip, which involves significant torsions of residues 380-388, leading to the insertion of Phe382 into the ATP-binding site, resulting in Abl1 inhibition.^33–36^ We tested our method on two mutations at position Phe382 in the DFG loop: F382L, which is expected to increase the inactive ground state population, ^37^ and F382V, which is expected to decrease it.^38^

**Figure 4:**
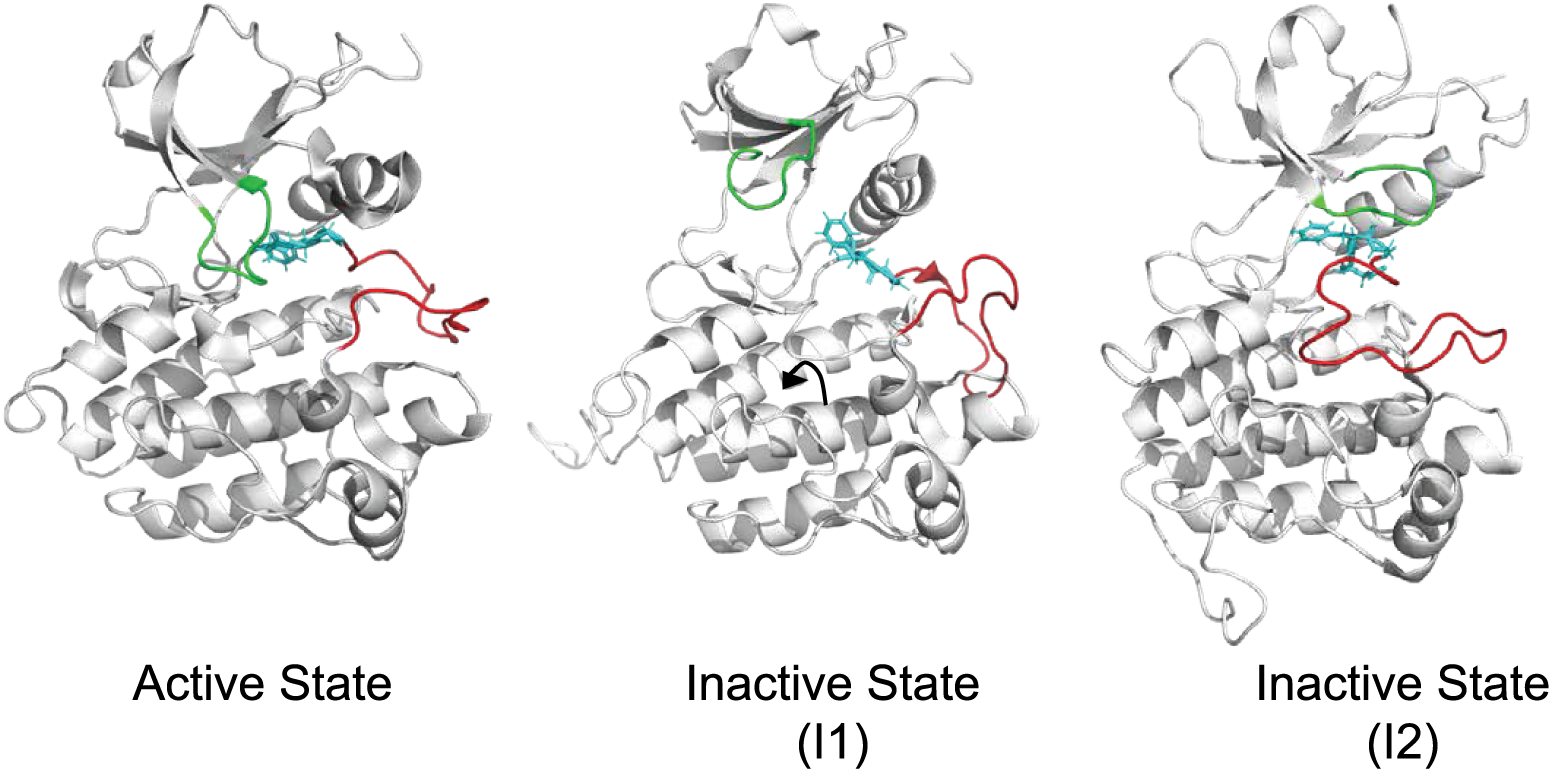
Abelson Kinase (Abl1) has one known active configuration (PDB 6XRX), and two distinct inactive configurations (PDB 6XR7, 6XRG).^39^ Regions of the kinase such as the P-loop (green), the DFG motif (cyan), and the A-loop (red) occupy different positions in each of these configurations.

In Figure 5A, the RMSF plots for the wild-type Abl1 protein show the fluctuation of each residue’s position throughout the ensemble of predicted structures. For the subsampling conditions 64:128 and 256:512, peaks around residue 150 indicate regions of high flexibility. These peaks suggest areas where the protein undergoes substantial conformational changes, which could be critical for its function and can motivate further investigations. To further validate this, we correlate this information with the pLDDT or TM-score confidence metrics, as well as any prior knowledge available about the protein, including known structures or stable states, experimental data, and information about protein function.

**Figure 5:**
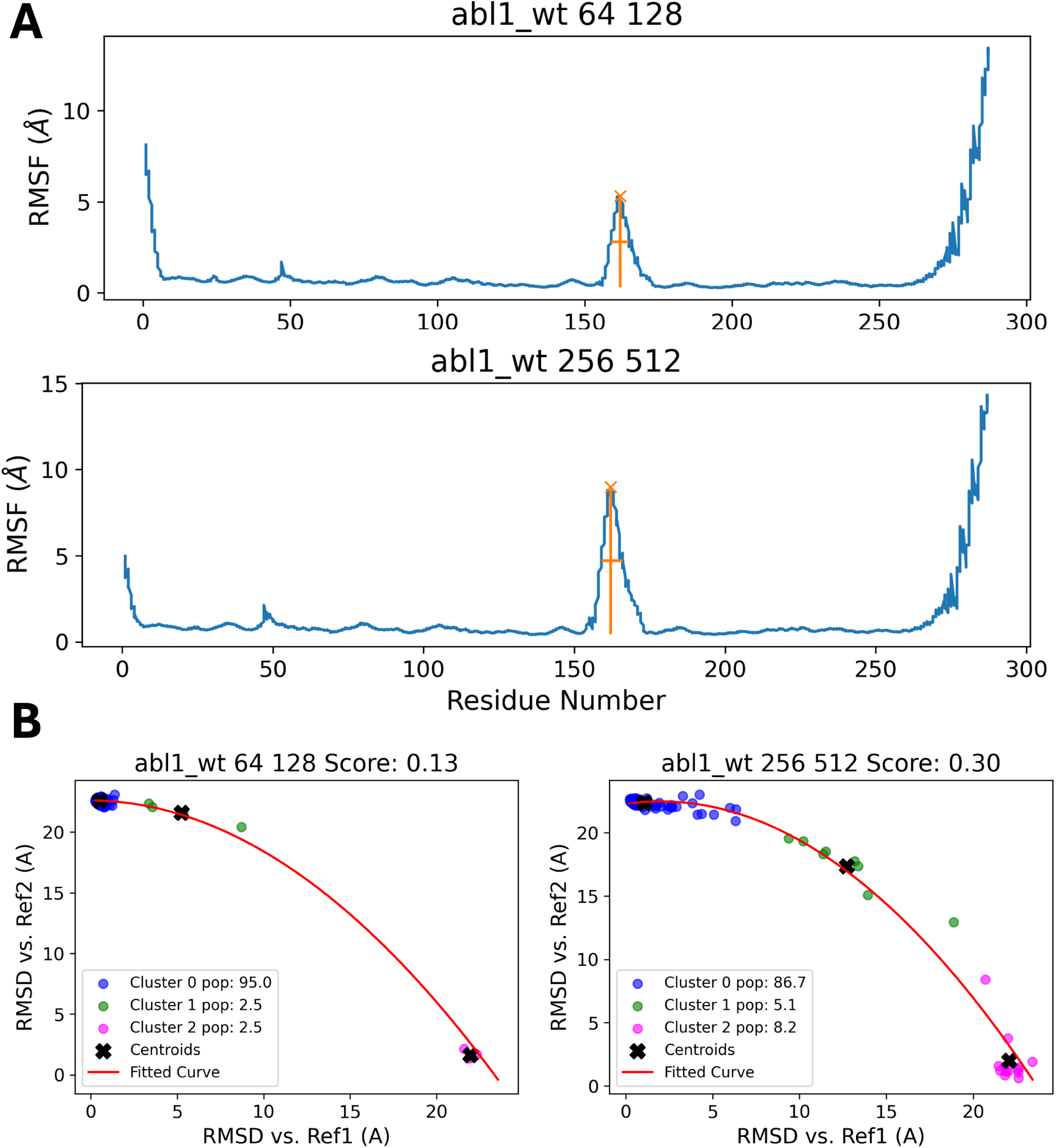
(A) RMSF plots of the wild-type Abl1 protein highlight residue flexibility, with a peak around residue 150 indicating regions of significant conformational change under 64:128 and 256:512 subsampling conditions. This area may play a key role in protein function. (B) RMSD values of different clusters from subsampling against two reference structures, representing active and inactive states. Cluster analysis reveals 95% of structures in the 64:128 condition belong to one conformation, while the 256:512 condition captures more diverse states. This suggests that 256:512 is optimal for predicting the effects of mutations on protein dynamics, as demonstrated by the Abl1 mutants F382L and F382V.

Figure 5B compares the RMSD values of different clusters, groups of similar conformations identified during the subsampling process, against two reference structures (Ref1 and Ref2). These reference structures are chosen from the ensemble to represent the active and inactive states of the protein, the two states between which we want to sample the trajectory. The scatter plots are color-coded to distinguish between the clusters, with centroids marked by black crosses and a fitted curve shown in red. The scores provided (e.g., 0.13 for Abl1 64:128) quantify the structural similarity between the clusters and the reference structures. The cluster analysis reveals that, for the Abl1 64:128 condition, Cluster 0 seems to be the predominant conformation (95% of the structures occupy this conformation). However, the 256:512 subsampling condition produces a larger variety of conformational states that lie between the two references and also shows a larger peak than that seen over the same residue range for the 64:128 subsampling in Figure 5A. This suggests that 256:512 could be an optimal set of subsampling parameters for this system, which we can use to predict the effects of point mutations on protein dynamics. Applying the same analysis to the Abl1 mutants F382L and F382V under similar conditions shows how specific mutations can have an effect on the relative populations of different conformational states.

As shown in Figure 6B, the population of cluster 0 is increased (86.7 to 87.3 percent) in Abl1 F382L, while the population of cluster 2 is decreased (8.9 to 3.8 percent) compared to Figures 5A and 5B). This shows that this mutation has an inactivating effect on Abl1, stabilizing the inactive conformation (cluster 0) over the active one (cluster 2). Interestingly, the population of cluster 1 is increased, which may be an indication of alternative stable states of the protein. In contrast, for Abl1 F382V, we observe the opposite effect; the population of cluster 0 is decreased (89.7 to 80.3 percent) and the population of cluster 2 is decreased (8.9 to 3.8 percent), while the population of the intermediate state cluster 1 is increased (4.4 to 15.9 percent). This is an example of how our method can be used to quickly and easily estimate the relative state populations given an ensemble of protein structures and compare the populations of the relative states to mutant variants.

**Figure 6:**
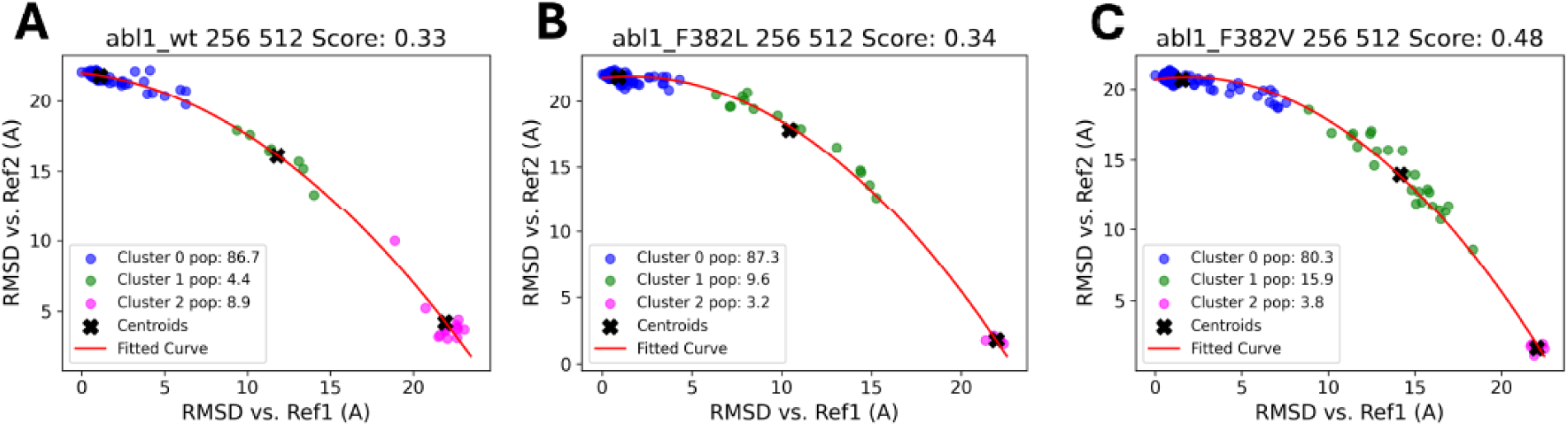
Comparison of the relative cluster populations across the Abl1 wildtype protein (A) and two mutants [Abl1 F382L shown in (B), Abl1 F382V shown in (C)], based on RMSD calculations for each structure in the ensemble plotted against structures from References 1 and 2.

### 3.2 Conformations of the LAT1 Protein

L-type amino acid transporter 1 (LAT1), also known as SLC7A5, is a heterodimeric protein that transports large neutral amino acids, thyroid hormones, and drugs across the plasma membrane^21^ (Figure 7). LAT1 is overexpressed in many types of tumors and mediates the transfer of drugs and hormones across the blood-brain barrier. ^41^ Over the years, interest in LAT1 has increased because this transporter is involved in important human diseases such as neurological disorders and cancer.^42^ Therefore, LAT1 has become an important pharmacological target together with other nutrient membrane transporters.^43^

**Figure 7:**
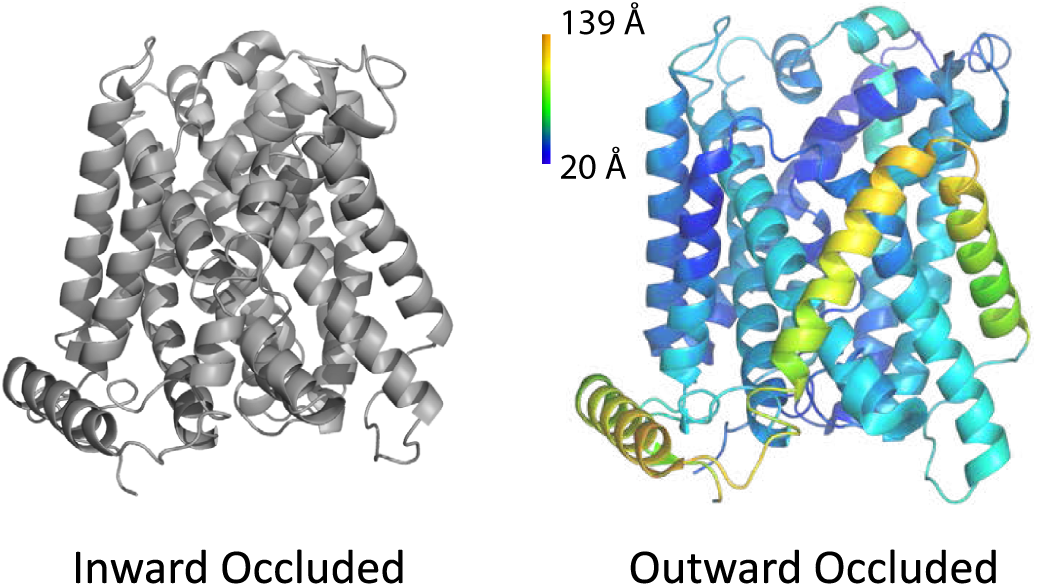
Cryo-EM structures of LAT1 show two different conformations: inward occluded (6IRS)^40^ and outward occluded (7DSK)^40^ color-coded on the figure on the right by per-residue RMSD between the outward occluded and inward occluded conformation. However, LAT1 is expected to have four different conformations to allow for successful transportation of substrates: an open conformation facing the extracellular side allowing for substrate binding (outward open); a closed conformation still facing the extracellular side (outward occluded); a closed conformation facing the intracellular side (inward occluded); and an open conformation facing intracellular side allowing for substrate release (inward open).

In Figure 8A, large RMSF values indicate flexible regions, while large pLDDT values represent regions of high confidence in the predicted structures. The color gradient shows the location of the residue in the protein sequence. RMSF vs. pLDDT plots for subsampling conditions 16:32 and 256:512 show that, although subsampling condition 16:32 might lead to a highly diverse structural ensemble (Figure 8B), it may also contain unphysical or unfolded states due to its low pLDDT score. In contrast, for subsampling condition 256:512, the average pLDDT is consistently high, showing that the prediction confidence score is large even in regions with large RMSF.

**Figure 8:**
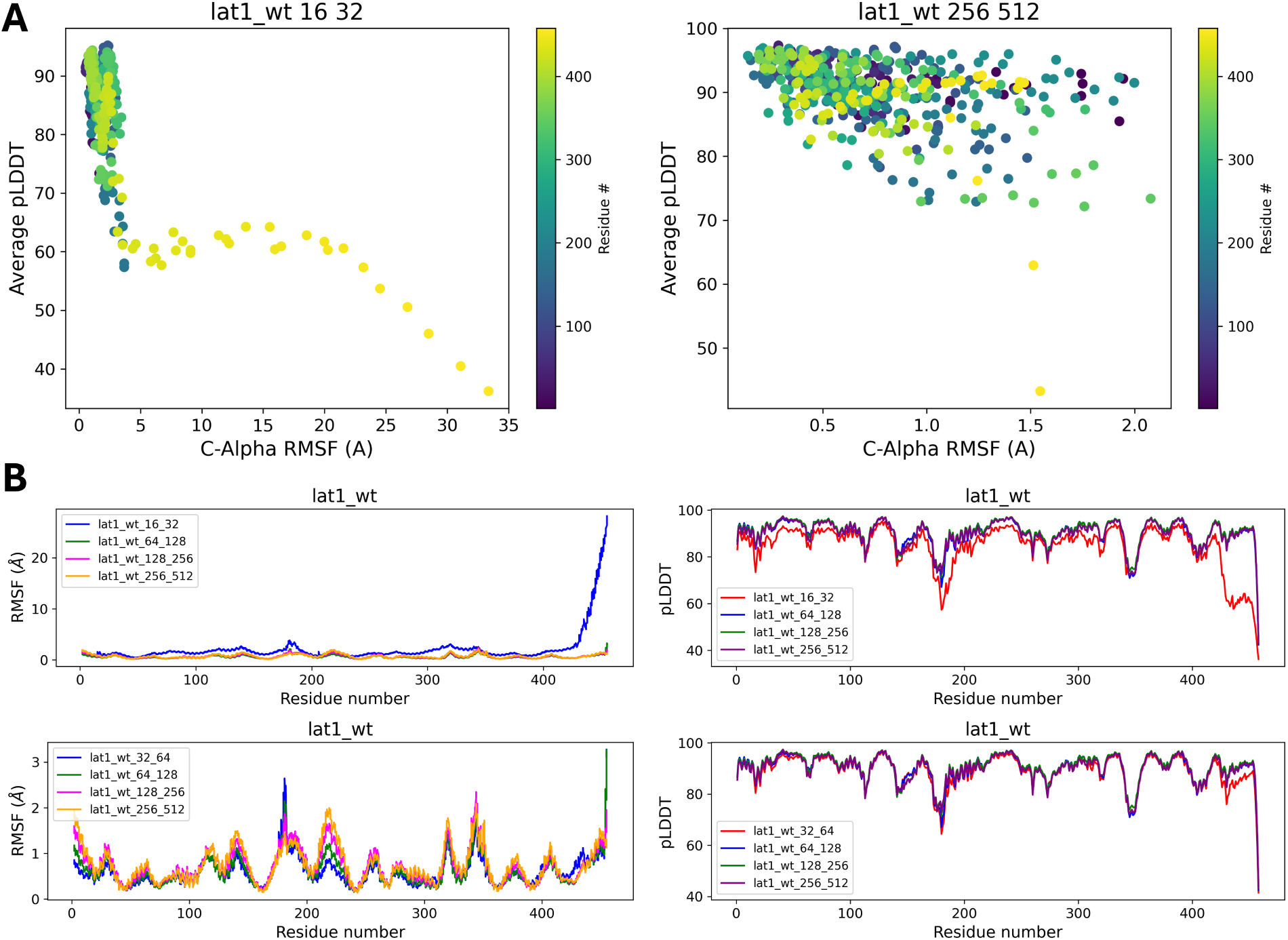
(A) pLDDT vs. RMSF plot of different subsampling parameters (16:32, 256:512) of the Lat1 kinase wild-type. The colorbar shows the index of each residue in the protein. (B) Plots of RMSF for every residue in the protein, showing peaks in highly dynamical regions of the protein.

Figure 8B shows 1D plots of the RMSF and pLDDT across all residues under different subsampling conditions (16:32, 64:128, 128:256, and 256:512). These plots consistently highlight flexible regions irrespective of sampling depth. Scores like 0.52 and 0.95 quantify the structural variability, with higher scores indicating more variability. The pLDDT plots show consistently high confidence across regions, even in dynamical regions of the protein such as residues 200-250, as shown in the RMSF plot.

In Figure 9, the visualization of cluster populations and centroids illustrates the structural diversity of the AlphaFold2-predicted structures under different subsampling parameters. We show that, compared to the subsampling condition 32:64, the subsampling condition 256:512 shows a more even distribution of the conformational state populations (cluster 0 has a 5% population for the 32:64 sampling condition as opposed to a 16.9% population for the 256:512 condition and cluster 2 has an 80.5% as opposed to a 63.1% population, respectively).

**Figure 9:**
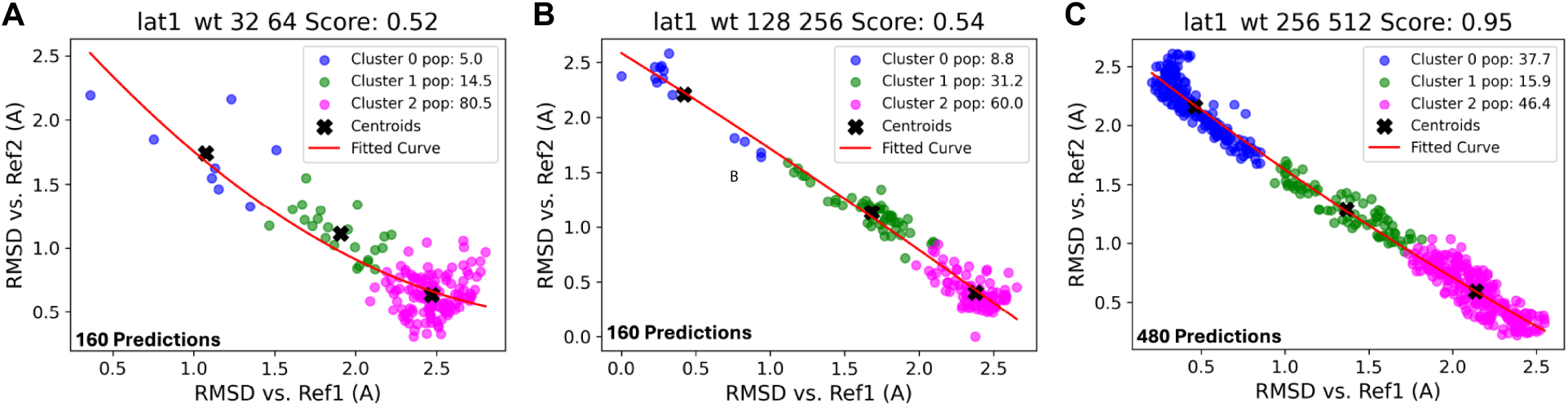
Comparison of the relative cluster populations across the LAT1 wildtype protein across different subsampling parameters (32:64, 128:256, 256:512, A-C respectively), based on RMSD calculations for each structure in the ensemble plotted against structures from References 1 and 2.

### 3.3 Conformations of the CCR5 Wild Type Protein

C-C chemokine receptor type 5 (CCR5) (Figure 10) is a G-Protein Coupled Receptor (GPCR) located on the surface of white blood cells that functions as a receptor for chemokines.^22^ Chemokine receptors have attracted substantial interest because they form portals into cells for the human immunodeficiency viruses (HIV-1 and HIV-2) and related simian or feline retroviruses, playing a vital role in immune surveillance and inflammation. ^22^ ^44^

**Figure 10:**
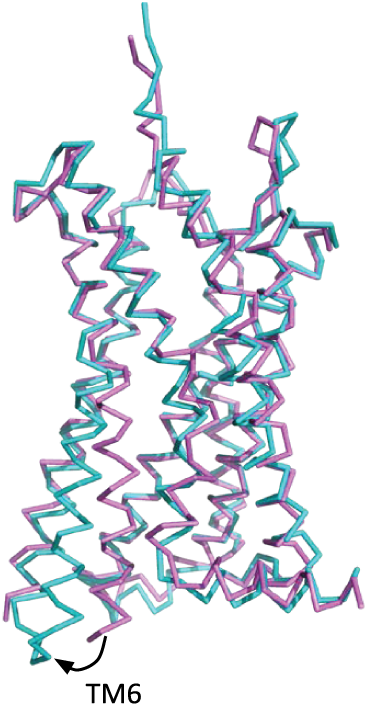
The intracellular portion of CCR5 changes conformation when transitioning from its active (PDB 7O7F) (teal)^45^ to its inactive conformation (PDB 5UIW) (magenta).^45^ The TM6 helix moves outward from the heptahelical bundle, causing further rearrangement of other secondary structures within the bundle.

The CCR5 wild-type protein was analyzed under various subsampling conditions to understand its conformational landscape. Figure 11A presents RMSF plots for CCR5 using the subsampling conditions 8:16 and 128:256, highlighting regions with significant fluctuations around residues 50, 100, 150, and 250. These peaks indicate flexible regions within the protein structure, which could be functionally important. We observe that there is consistency between these two subsampling conditions in predicting that, around residues 50 and 200-250, there must be highly dynamical regions.

**Figure 11:**
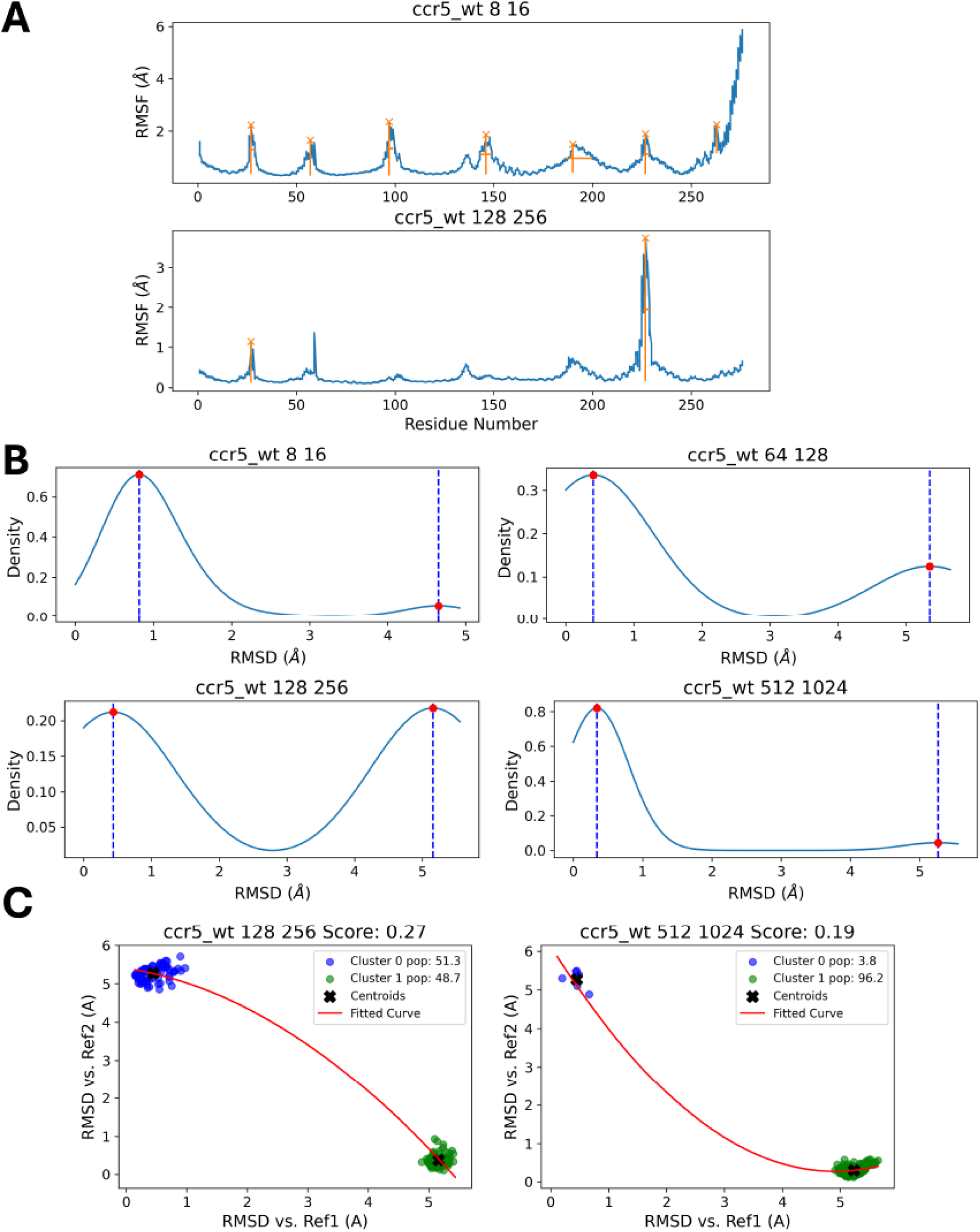
(A) RMSF plots for CCR5 using the subsampling conditions 8:16 and 128:256, highlighting flexible regions around residues 50, 100, 150, and 250, which signal highly dynamic areas. (B) Density plots of RMSD values, indicating multiple conformational states. (C) RMSD scatter plots for subsampling conditions 128:256 and 512:1024, comparing clusters to the structures observed in References 1 and 2, with similarity scores (0.27 and 0.19), reflecting varying degrees of structural similarity.

Figure 11B presents density plots that display the distribution of RMSD values for different subsampling conditions. The presence of multiple peaks in these density plots suggests multiple conformational states of the protein. In Figure 11C, the RMSD scatter plots for the CCR5 wild type protein (under conditions 128:256 and 512:1024) show how different clusters relate to the reference structures. The similarity scores (0.27 and 0.19) suggest varying degrees of similarity between the clusters and the references, with lower scores indicating greater structural similarity.

Together, these examples illustrate the versatility of our tool in handling proteins with different degrees of complexity and different types of conformational ensembles.

## 4 Conclusions

FastConformation offers a versatile and user-friendly Python-based solution for predicting and analyzing protein conformational ensembles using subsampled AlphaFold 2 predictions. The application provides all necessary functionalities for predicting and analyzing protein conformational ensembles in one place, enabling users to start from an amino acid sequence and achieve protein conformation prediction and analysis in just a few hours on their local machine. This is substantially faster than traditional molecular dynamics simulations, making it applicable for high-throughput screening. By integrating MSA generation, AF2 predictions, and interactive visualization, our tool allows users to explore alternative protein conformations and assess the impact of mutations on protein conformational ensembles. The application’s GUI makes it accessible to users without programming knowledge, broadening its utility in various fields including biophysics, drug discovery, and protein engineering. Our validation on the Abl1 kinase, LAT1 transporter, and CCR5 receptor demonstrates the robustness and versatility of this approach. We expect FastConformation to be instrumental in bridging the gap between protein sequence, structure, and function, by providing insights into protein conformations starting from a target sequence. FastConformation holds promise in advancing personalized medicine and therapeutic development.

## Associated Content

To accompany this manuscript and the FastConformation code, we have designed a FastConformation GitHub page, where installation directions and examples can be found: https://github.com/GMdSilva/FastConformation. We have also illustrated the use of the package through this YouTube video: https://www.youtube.com/watch?v=a7_y4lxic8w.

## Acknowledgement

The authors thank F. Marty Ytreberg, Jagdish S. Patel, Marcelo D. Pol^eto, Kyle Lam, Gerwald Jogl, Lillian Chong, Patrycja Dubielecka-Szczerba, and George Lisi for insightful conversations and support. G.M.d.S. was supported by a Blavatnik Family Fellowship award. G.M.d.S. and B.M.R. were supported in part by the National Science Foundation Grant No. 2027108 and CTMC CAREER Award 2046744. I.V. was supported by the NIGMS T32 Training Grant in Interdisciplinary Pharmacological Sciences Award Number T32GM139793. This research was conducted using computational resources and services at the Center for Computation and Visualization, Brown University.

